# Formulation and Characterization of Primary Tissue-Derived Microfluidic Droplet-Engineered Organoids

**DOI:** 10.1101/2025.07.10.664241

**Authors:** Wanlong Wang, Yongde Cai, Xiaoyong Dai, Haowei Yang, Davit Khutsishvili, Jiawei Li, Yu Zhu, Jin Wang, Xiu Yan, Zitian Wang, Shaohua Ma

**Author notes:** These authors contributed equally to this work (Wanlong Wang, Yongde Cai).

## Abstract

Organoid technology offers a powerful platform for modeling human tissues, studying disease mechanisms, and developing personalized therapies. However, widespread clinical application is hindered by challenges in scalability, reproducibility, and the handling of ultra-small tissue samples typical of clinical biopsies. Here, we present a comprehensive and automated protocol for the formulation and characterization of microfluidic droplet-engineered organoids (DEOs) derived from primary tissue samples. This protocol integrates 3 key stages: (1) Extraction and purification of viable primary cells from ultra-small tissue specimens using the small-Tissue Extraction Device (sTED); (2) High-throughput fabrication of uniform cell-laden microspheres using an integrated microfluidic bioprinter (OrgFab), capable of generating over 100 organoids from just 10 μL of bioink; and (3) Rapid organoid characterization using lamination-based processing for single-cell analysis while preserving spatial context.

The automated workflow minimizes manual intervention, reducing variability and enhancing reproducibility, making it suitable for high-throughput applications such as drug screening and disease modeling. Our method allows for the generation of patient-derived organoids that closely mimic the native tissue microenvironment, including diverse cell types and structural features, within a significantly reduced timeframe. This 3 stage protocol enhances the use of organoids in personalized medicine by enabling the rapid assessment of drug efficacy in patient-specific models. Here, the integration of advanced techniques supports existing organoid protocols, providing a valuable resource for researchers and clinicians seeking to improve patient outcomes through more efficient and precise organoid-based applications.

## Introduction

Cancerous tumors remain a leading global health threat, with persistently high incidence and mortality rates. Their intrinsic characteristics, such as tumor heterogeneity and dynamic evolution, pose significant challenges for traditional treatments and even genetic sequencing-based precision medicine^1^. Rapid and extensive screening of tumors using patient-derived genetic and epigenetic living models, particularly those generated from ultra-small biopsy samples, is crucial for identifying the most effective treatments and improving patient outcomes^2,3^.

Three-dimensional (3D) organoid models have emerged as powerful tools for replicating the architecture and function of original tissues, providing invaluable platforms for disease modeling^4^, drug screening^5^, and personalized medicine^6,7^. Organoids derived from patient tumors, known as patient-derived organoids (PDOs), retain the genetic and histological features of the original tumor, enabling more accurate predictions of therapeutic responses^2,3,8,9^. However, the widespread clinical application of organoid technology faces significant challenges. Traditional methods for organoid culture are labor-intensive, time-consuming, and suffer from issues related to scalability and reproducibility^10–14^. Moreover, handling ultra-small tissue samples, which are common in clinical biopsies, remains particularly problematic due to limited cell numbers and difficulties in maintaining viability^15,16^.

Recent advancements in automated cell extraction, microfluidic organoid formulation, and spatial transcriptomics offer promising solutions to these challenges^3,17–23^. Automated tissue dissociation devices enable efficient and reproducible extraction of viable cells from small tissue samples^24,25^. Microfluidic technologies facilitate the high-throughput formation of uniform organoid microspheres, enhancing scalability and consistency^26^. While single-cell sequencing analysis is rapidly advancing, revealing more about cell-cell communication^27–29^, the integration of spatial transcriptomics enables the comprehensive characterization of organoids at the single-cell level while preserving spatial context, providing insights into cellular heterogeneity and tumor microenvironment interactions^8,30^.

This protocol presents a comprehensive, high-throughput system for treating, generating, and characterizing patient-derived organoids from ultra-small tissue samples. It integrates automated cell extraction, microfluidic droplet-based organoid printing, and lamination-based spatial transcriptomics. Streamlining organoid workflow presents an opportunity to accelerate the clinical translation of organoid technology, enabling applications such as personalized therapy selection in a shorter timeframe.

## Development of the Protocol

This protocol builds upon previous advancements in primary tissue cell extraction, organoid microsphere formulation and printing^26^, and organoid profiling^8,9^. Our focus is on primary tissue or patient-derived organoids, emphasizing high-throughput formation and automated processing to enhance scalability and reproducibility.

The protocol includes three key stages (Figure 1):

1. Extraction and Purification of Primary Cells from Tissue Samples (Figure 2)
2. Formulation and Bioprinting of Cell-Laden Microspheres (Figure 3)
3. Lamination of Organoids for Rapid Characterization (Figure 4)

**Figure 1.**
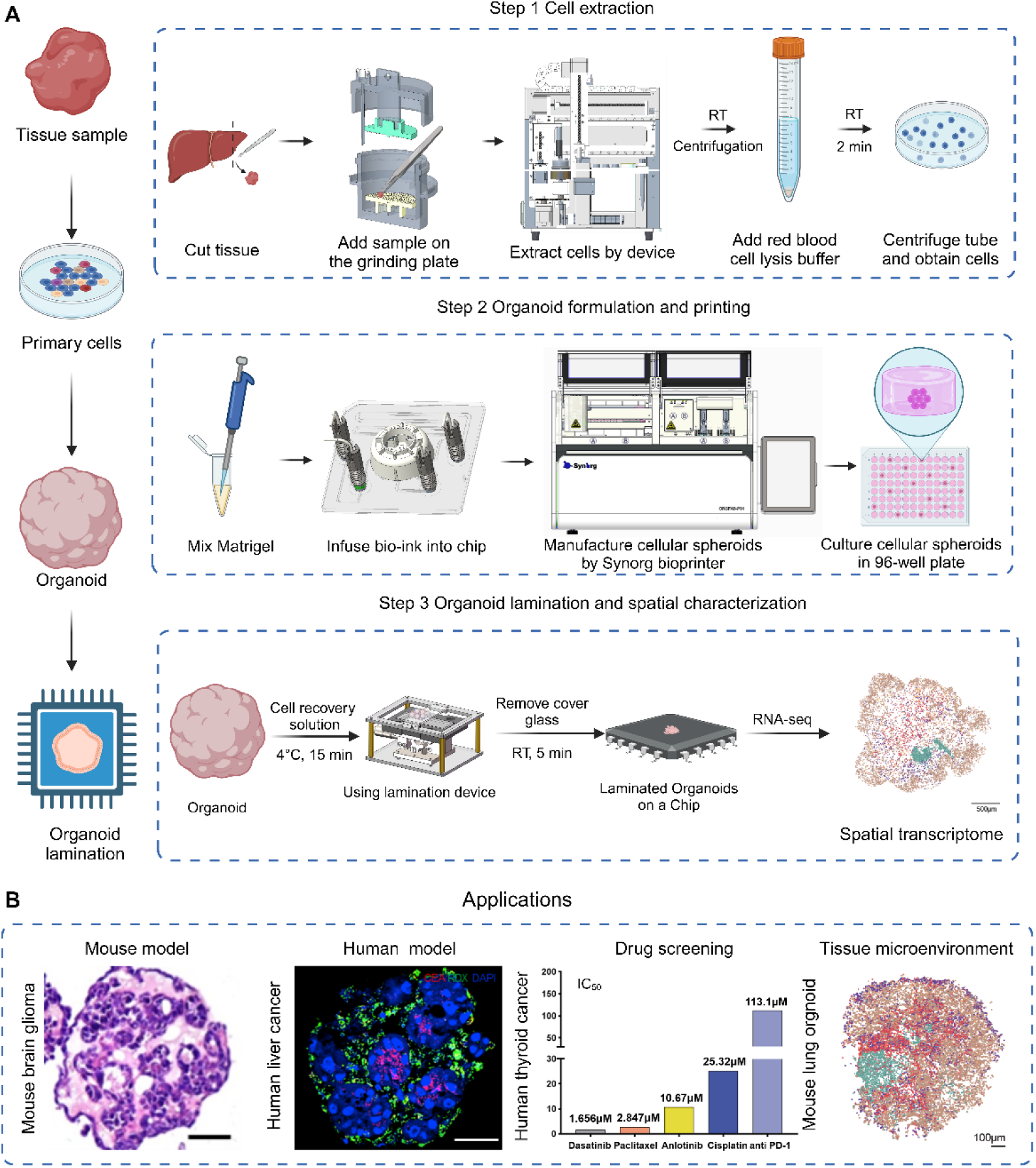
Workflow to formulate and characterize microfluidic droplet-engineered organoids (DEOs) A) Step 1. Extract cells from tissue, including using forceps and scalpels to cut tissue into small pieces, grinding tissue cuts using the automated small-Tissue Extraction Device (sTED), red blood cell lysis, and centrifuging cell suspension to obtain cell pellets. Step 2. Formulate and print DEOs using extracted cells from Step 1, including suspending cells in Matrigel to create bioink, infusing bioink to sampling chip, and installing chip to OrgFab printer to formulate and print DEOs into 96-well plates. Details are described in Box 2. Step 3. DEO profiling by laminating 3D organoid cells into 2D mono-layer distribution. It includes Matrigel removal by matrix digestion, laminating organoids on a spatial transcriptome chip using a lamination device, and spatial sequencing profiling. B) Application examples of **DEOs**. From left to right: HE staining of mouse brain glioma organoids, fluorescent staining of human liver cancer organoids (red: CEA, green: RDX, blue: DAPI), anticancer drug screening of human thyroid cancer organoids (IC50, the drug concentration at which anticancer drugs inhibit tumor cell activity by 50%), visualization of cell annotation in spatial transcriptomics of a primary lung organoid, scale bar, 100μm.

**Figure 2.**
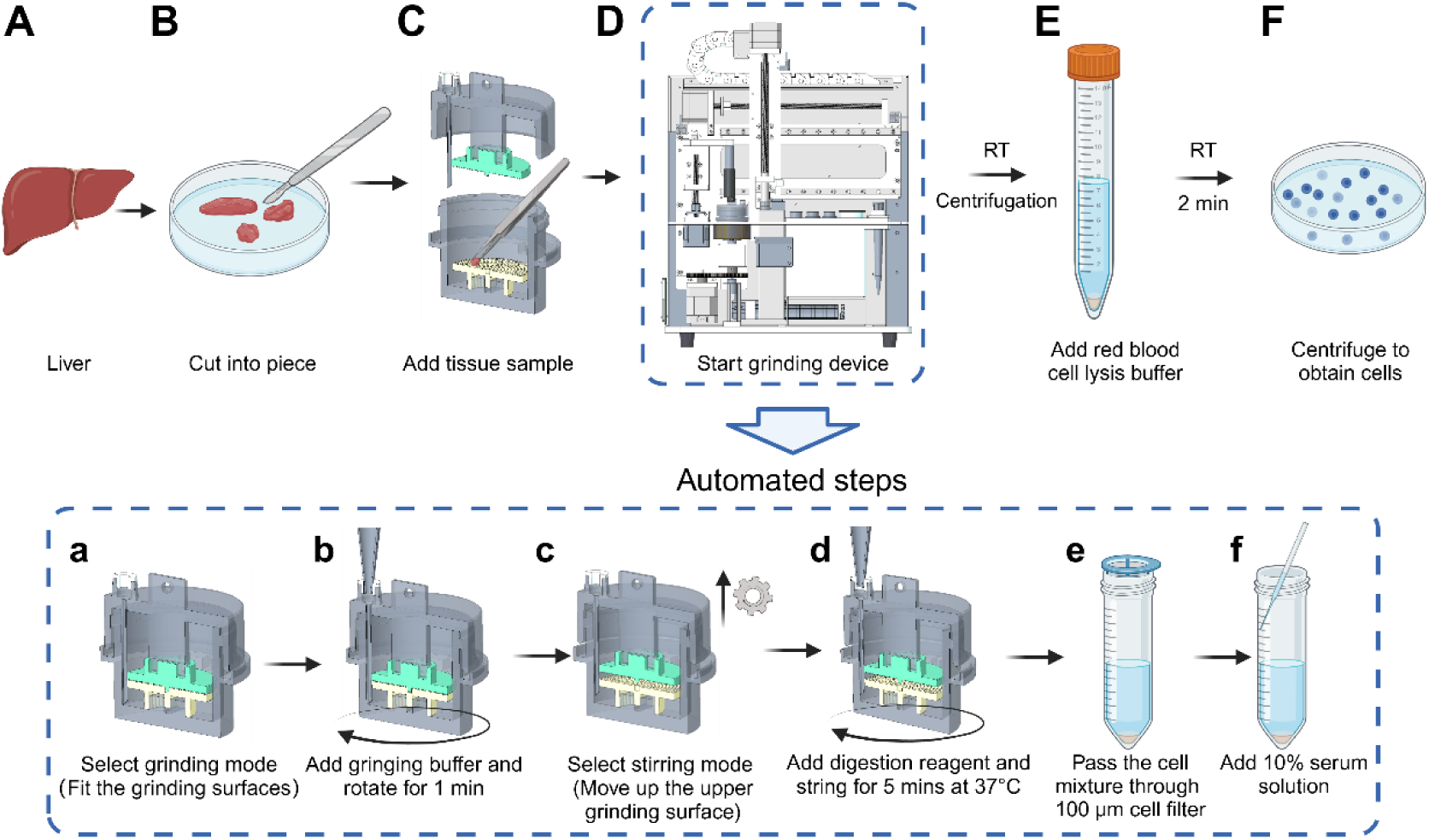
Workflow to extract cells from primary tissues. A. Select liver tissue from BALB/c mouse
B. Use forceps and sterilized scalpels to cut the primary tissue sample on ice into small fragments (approximately < 1mm in size)
C. Open the grinding chamber, and add the tissue fragments with forceps
D. Run the grinding device, which includes 6 steps (a-f):

a. Select the grinding mode, and lower the upper grinding plate until the grinding surfaces fuse together
b. Add the grinding buffer, set the lower grinding plate rotating at 60r/min, and start the motor to run for one minute
c. Select the stirring mode and lift the upper grinding plate for 10 mm to separate the grinding plates
d. Add 10× tissue digestion reagent, and rotate the lower grinding plate for 5 min at 60r/min and 37 °C
e. Pass the cell suspension through a 100 μm cell filter
f. Add an equal volume of digestion termination reagent, which contains 10% serum, to the cell suspension
E. Centrifuge the cell suspension, remove the supernatant, add 2 mL of red blood cell lysis solution to the cell pellet, pipette the mixture, and incubate it at room temperature for 2 min
F. Re-centrifuge the cell suspension, remove the supernatant, add 2 mL culture medium to the cell pellet, and pipette the mixture to obtain primary cell suspension

**Figure 3.**
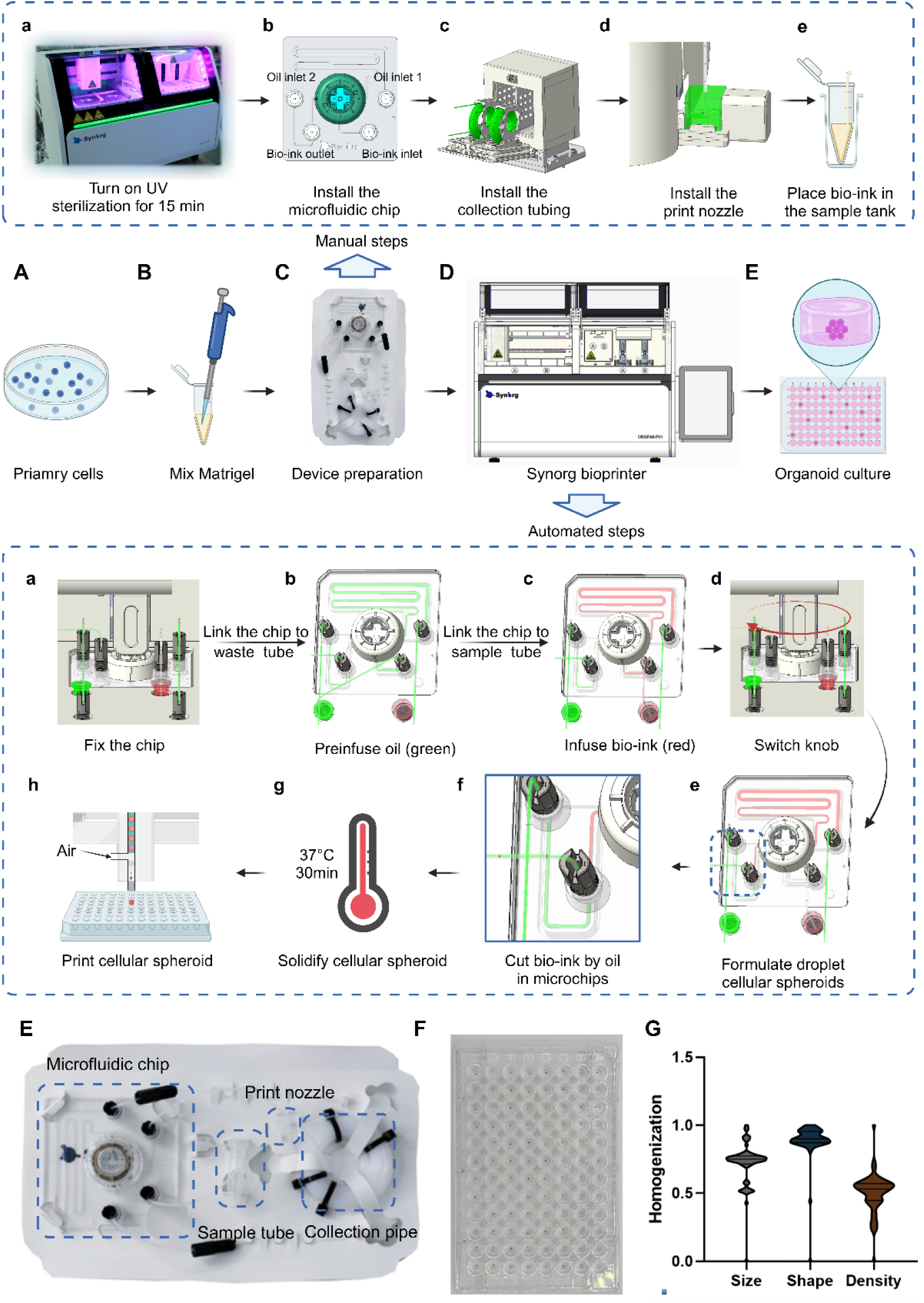
Workflow to formulate and print organoid precursor, i.e. cellular spheroids, using the OrgFab printer. A. Centrifuge the cell suspension and remove the supernatant to obtain a cell pellet
B. Use a pre-cooled pipette tip to aspirate Matrigel on ice, pipette it to the EP tube, and mix with cells by gently pipetting
C. Manual preparation steps before using the OrgFab printer (a-e):

a. Turn on UV sterilization for 20 min
b. Install the microfluidic chip to the semiconductor cooling platform, and connect the inlet/outlet ports to corresponding fluids, including two oil inlet ports, one bioink inlet port, and one droplet outlet port
c. Insert bioink inlet tubing into the bioink EP tube, aspirate oil out of the channel from the front right (Oil inlet 1) to port aspirate bioink with flow rates of 250uL/min from the tube, and infuse the channel
d. Install the print head, and connect one end of the print head to the collection/incubation tubing and the other end to the high-pressure gas. The third port is placed facing straight to the recipient substrate, e. g. 96-well plate
e. Place the EP tube with bioink in the sample tank and insert the bioink tube
D. OrgFab printer operation, which includes 8 automated steps (a-h):

a. Move down the mechanical turntable to fix the chip on the semiconductor cooling platform
b. Insert the chip bioink inlet tubing into the EP tube for waste collection, and pump oil into the chip channels with flow rates of 312.5uL/min via the two oil inlets to repel air in the channel. Afterward, the chip channels are filled with oil. All fluids are conducted via PTFE tubing
c. Insert bioink inlet tubing into the bioink EP tube, aspirate bioink from the tube, and infuse the channel, meanwhile, perfuse oil out of the channel segment connecting to the front right (inlet) port
d. Rotate the mechanical turntable (knob) to change the chip channel tandem, which connects the bioink channel to the front left port for the bioink outlet
e. Pump oil via the two oil inlets, with flow rates of 312.5uL/min in Oil inlet 2 and 25uL/min in Oil inlet 1
f. Shear bioink flow into uniform droplets by oil co-flow, forming structural templates of organoid precursors (or cellular spheroids)
g. Stop pumping oil toward the two inlet channels, store the cell-laden droplets in the collection tubing, and solidify them at 37°C for 30 min
h. Pump oil from Oil inlet 1 at 12.5uL/min and Oil inlet 2 at 25uL/min, use a camera and visual detection algorithm to control the instantaneous opening of the high-pressure gas toward the print head. Each air beam firing enables one shot of gelled droplet, i.e. organoid precursor/cellular spheroid, into the 96-well plate, with each well containing one or a certain number of spheroid
E. Culture organoid precursors/cellular spheroids in a 96-well plate
F. Picture of the consumable microfluidic chip module
G. Picture of organoid precursors after printing in a 96-well plate
H. Violin plot of area and roundness of printed organoids.

**Figure 4.**
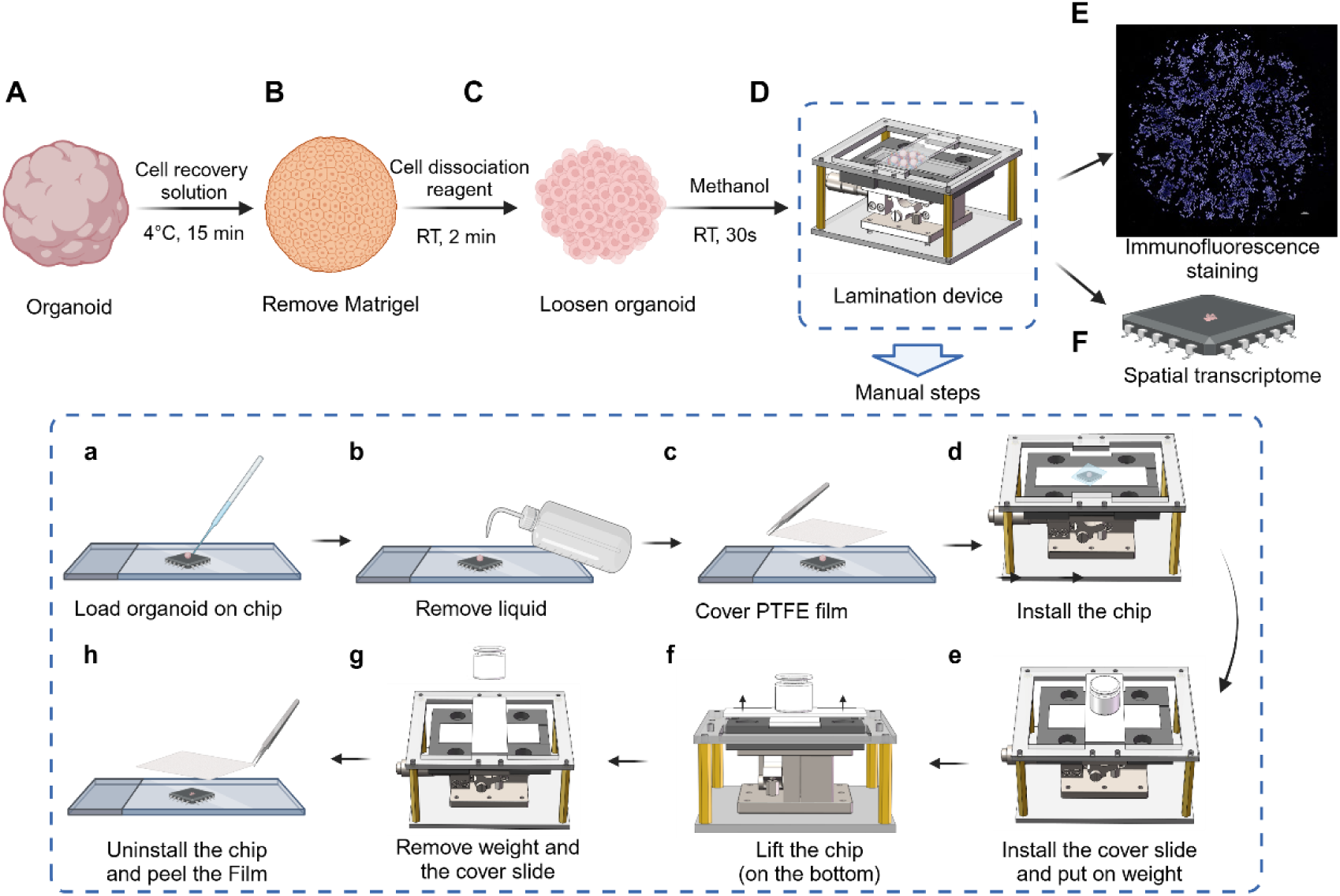
Workflow to laminate organoids on the chip. A. Select and pick out organoids using a Pasteur pipette under the guidance of a microscope
B. Clean the culture medium by repeated washing with 1× SSC buffer, add cell recovery solution for the Organoid, and incubate it at 4°C for 15min
C. Wash the cell recovery solution for Organoid with 1× SSC buffer, add cell dissociation reagent, and incubate at room temperature for 2min
D. Use the lamination device to create 2D cell distribution, which includes 8 manual steps (a-h):

a. Use a Pasteur pipette to load the organoid on the chip
b. Use a gas cylinder to blow and dry the liquid around the organoid
c. Use tweezers to cover the chip with a piece of PTFE film
d. Install the chip on the lamination device
e. Place a glass slide on the laminating device and place a weight on top of the glass slide
f. Switch the knob to lift the chip until it touches the glass slide installed on the top
g. Gently and vertically remove the weight and then the glass slide
h. Uninstall the chip from the lamination device, heat the chip at 37°C for 5 min, and use tweezers to remove the PTFE film
E. Immunofluorescence staining of a laminated organoid, showing DAPI staining of the laminated cellular spheroid. Scale bar=100μm
F. Laminated organoids on a spatial transcriptomic chip, enabling sequencing and resolution of cell types and cellular interactions

All three steps are performed using automated devices (OrgFab, Synorg Biotechnology, Shenzhen). Manual sample transport is required between stages and devices.

### Stage 1: Obtaining Primary Cells from Tissues

The acquisition of viable primary cells is critical for constructing accurate patient tissue-derived organoid models. Traditional methods often involve manual grinding of tissue followed by enzymatic digestion^31–34^. However, manual techniques are inherently variable, as the degree of tissue fragmentation and operator proficiency can significantly affect cell viability and yield^14,35^. For instance, inconsistencies in tissue preparation can impact the establishment and growth of organoids^36^. Moreover, specialized tissues like the liver require perfusion-based extraction methods, which are complex and difficult to standardize^37^.

To address these challenges, we employ the automated small-Tissue Extraction Device that standardizes tissue processing (Figure 2D), minimizing operator-dependent variability and enhancing the consistency of primary cell isolation.

### Stage 2: Constructing Primary Cell-Based *In Vitro* 3D Cell Microspheres

The streamlined, automated process not only improves the spatial precision and reproducibility of organoid precursor fabrication but also enhances scalability for high-throughput applications in drug screening and disease modeling^38–40^.

At this stage, we focus on the automated fabrication of primary cell-based in vitro 3D microspheres using an integrated microfluidic bioprinting system. This approach addresses the challenges of scalability, consistency, and handling ultra-small sample sizes in organoid production. Building upon advancements reported in our previous work on the OrgFab printer^26^, primary cells are combined with Matrigel to form a bioink, which is then processed using a microfluidic technique employing oil-phase shearing to generate uniform, multi-cell-laden droplets. These droplets undergo online Matrigel solidification, ensuring immediate stabilization of the organoid precursors. Subsequently, a 3D bioprinting nozzle precisely deposits the solidified microspheres onto multi-well plates, our target substrates for further culture (Figure 3D). The integration of microfluidics and bioprinting technologies allows for high spatial precision in droplet formation and placement, significantly improving the reproducibility of organoid fabrication.

The streamlined process not only enhances spatial precision but also improves scalability for high-throughput applications in drug screening and disease modeling. The OrgFab system is capable of handling ultra-small sample volumes down to 5 μL, producing more than 100 organoid precursors from just 10 μL of bioink. For example, this is valuable for needle biopsy samples.

The fully automated workflow minimizes manual intervention, reducing variability and enhancing reproducibility. The OrgFab printer integrates a computer vision-based bioprinting module, which enables accurate detection and deposition of organoid precursors into multi-well plates. This precision facilitates downstream high-throughput assays and ensures consistency across experiments without requiring an operator. Additionally, the high-density microfluidic droplet encapsulation used in this method has been shown to accelerate organoid maturation due to increased paracrine signaling among densely packed cells^26^. This results in more rapid organoid development and the potential for earlier experimental readouts, which is advantageous in time-sensitive research and clinical contexts. By integrating these techniques, our protocol tries to address the key limitations of existing methods, such as low throughput and limited automation^38,40–42^. It provides an integrated platform for the controllable modeling of organoids, with promising developments for personalized medicine, drug discovery, and disease modeling.

### Stage 3: Laminating Organoids for Rapid Characterization

Traditional methods for organoid characterization, such as sectioning, immunostaining, and even more recent single-cell sequencing, are labor-intensive and can result in cell loss and incomplete spatial information. Recent advancements in spatial transcriptomics have enabled high-resolution mapping of gene expression within tissues and organoids^43–45^. However, applying these techniques to primary tissue-derived organoids remains challenging due to their small size and low cell count, which limits the feasibility of tissue sectioning as a pre-processing step before profiling.

Based on these limitations and advancements, we developed a lamination device that compresses organoids into a single-cell-layer morphology without disrupting cellular architecture (Figure 4, Figure 7). This lamination technique allows projecting 3D organoid spheres onto a 2D plane, preserving spatial relationships between cells. By integrating lamination, which enables spatial transcriptomics profiling, our protocol facilitates the comprehensive characterization of organoids at the single-cell level, followed by immunofluorescence staining. This approach is particularly valuable for understanding cellular heterogeneity and interactions within tumor organoids.

## Applications of the Methods

This three-stage protocol establishes a controlled platform for the generation and high-throughput characterization of primary tissue-derived organoids from both healthy and cancerous tissues, which can be derived from human patients or animal models (e.g., murine). Various stages of this protocol have been modeled for multiple cancer types, including lung, liver, thyroid, colorectal, breast, gastric, and cervical cancer, as well as brain glioma, chocolate cyst, and skin keloid (Supplementary Figure 3), making it highly versatile for preclinical research^3,38^.

By standardizing the extraction of viable primary cells and integrating a microfluidic bioprinting system to fabricate uniform, cell-laden 3D microspheres, this method enables the construction of rapidly reproducible *in vivo* tumor models, such as primary mouse liver and lung cancer organoids, for patient-specific drug screening, resistance mechanism studies, tumor and immune microenvironment modeling, metastasis driver identification, and novel drug mechanism research - including toxicity, metabolism, and biomarker discovery^8,26^.

Furthermore, by employing a lamination device that converts complex 3D organoid spheres into a single-cell-layer without compromising cellular architecture, the protocol facilitates rapid spatial transcriptomic and immunofluorescence analyses, vital for personalized drug screening and in-depth analysis of tumor heterogeneity and therapeutic responses in patient-derived models^9^.

### Comparison with other Methods

Several approaches have been proposed to address the challenges of variability, scalability, and heterogeneity in organoid research. While these methods have advanced automation, throughput, or analysis, they often focus on specific aspects of the organoid workflow or are limited to particular cell types, such as pluripotent stem cell (PSC)-derived organoids generated through controlled cascade differentiation.

Harrison et al. developed a method to produce liver organoids from human PSCs using small molecules that mimic embryonic liver development without relying on an extracellular matrix. Their approach bypasses 2D patterning and conventional growth factors, yielding organoids with vascular structures and Kupffer cells that function in vivo over time; however, it is limited to PSC-derived liver organoids and may not translate to primary tissues or other organoid types^46^.

Schuster et al. created an automated microfluidic platform for culturing 3D tumor organoids and performing combinatorial drug screening. Their system supports high-throughput culture and real-time analysis, as demonstrated with human pancreatic tumor organoids where drug responses varied among patient samples. This platform does not address primary tissue processing and limits the system’s capacity to produce uniform, large organoids with high cellular diversity^41^.

Beghin et al. introduced an automated 3D imaging system that combines a single-objective light-sheet microscope with disposable microfabricated “JeWells” chips to image live organoids at a rate of about 300 per hour. The system allows phenotypic quantification at multiple scales but does not include methods for organoid generation or processing of limited tissue samples. More importantly, optical methods are restricted by light scattering, limiting their ability to penetrate deep into tissues and acquire detailed cellular information.^47^

Bozano et al. developed an automated 3D high-content screening platform in a 384-well format that integrates robotic liquid handling with confocal imaging for drug screening of 3D organoid cultures. Their work indicates that automation can enhance precision and throughput compared to manual methods, with confocal imaging detecting drug-induced changes more sensitively. This system is focused on screening pre-formed organoids^48^.

Liu et al. designed a droplet microfluidic system to form hybrid hydrogel capsules via interfacial complexation of oppositely charged polymers. This all-in-water approach enables continuous production of islet organoids from human induced pluripotent stem cells that exhibit glucose-responsive insulin secretion, though the method is specific to hiPSC-derived islet organoids and may not suit primary patient tissues^49^.

Additional methods include a spatially and optically tailored 3D printing process by Sanchez Noriega et al., a digital microfluidic platform for single-cell isolation (DISCO) by Lamanna et al., and a high-throughput approach for isolating circulating tumor cell clusters by Boya et al. These techniques improve specific aspects of microfluidics and cell handling but do not offer an integrated solution for automated organoid fabrication and characterization from ultra-small tissue samples^50–52^.

Mathur et al. developed Combi-Seq, a microfluidic workflow that uses deterministic barcoding in single-cell droplets for multiplexed transcriptome profiling of hundreds of drug combinations in picoliter droplets. While this method can predict drug interactions from the minimal sample input, it requires pre-prepared cell suspensions and focuses on microfluidic-enabled drug combinations and transcriptomic analysis rather than on organoid fabrication^53^.

In contrast, our protocol offers a comprehensive solution by integrating automated cell extraction optimized for ultra-small primary tissue samples through sTED, ensuring high cell viability and yield from limited biopsy material (**Stage 1**). The microfluidic droplet-based organoid printer enables high-throughput, uniform fabrication (**Stage 2**), while the lamination-based spatial transcriptomics technique allows for advanced single-cell characterization while preserving spatial context (**Stage 3**). Unlike methods that target specific organoid types or rely on stem cell-derived models, our protocol applies to various primary tissues, increasing its translational potential for clinical applications where sample size is limited.

### Limitations

While our protocol offers advantages from using primary cells, such as cellular diversity, natural self-organization, and reduced dependence on exogenous growth factors, it can be improved. For instance, the method currently relies on an extracellular matrix (ECM) component, such as Matrigel, which necessitates the use of synthetic hydrogels that can reduce batch-to-batch variability and may require adaptation for certain cell types. Additionally, although we have streamlined many steps, manual sample transport between stages and devices is still required, which may introduce minor variability. Although the protocol is detailed, operator experience with tissue engineering yields a better outcome, as some variables will be adjusted in a tissue type-dependent manner. In addition, the process of this protocol is dependent on specific equipment. To ensure that potential users can utilize this protocol, we have added the steps for manually constructing a microfluidic platform (SI). Future iterations aim to further automate these transitions to boost reproducibility and scalability.

Moreover, our lamination-based spatial transcriptomics approach effectively preserves key spatial information while expediting analysis, yet the process of compressing organoids into a planar format can sometimes diminish the full complexity of their three-dimensional morphology, particularly for cell types with intricate architectures. We are actively exploring computational algorithms and imaging enhancements to better recapture the original three-dimensional context. Furthermore, our protocol is optimized for primary cell cues, making it well suited for generating organoids that closely mirror native tissue function; however, this means it is not immediately compatible with induced pluripotent stem cell (iPSC) or embryonic stem cell (ESC) models, which typically require additional growth factor support. Finally, although the resulting organoids are produced with high consistency and throughput, they tend to be smaller and somewhat simpler in structure compared to traditional 3D bioprinting tissue models, a trade-off for achieving rapid, scale-out and reproducible production. Overall, these limitations provide clear targets for future improvements.

### Experimental Design

In order to automate the process from primary tissue to organoid manufacturing and characterization, we used three devices to achieve cell extraction, organoid manufacturing, and characterization automation. The total process took 3 hours (main step) + 7-14 days (culture).

#### Box 1 | Extract cells from primary tissues Timing 30 min Procedure

**▲CRITICAL** The cell extraction process must be performed within a sterile laminar flow hood. Prior to commencing the procedure, the equipment should be sterilized with ultraviolet light.

1. Take out one 60 mm diameter Petri dish, add 2 mL of DPBS (add 2% P/S), and place the tissue block to be processed into the dish.
2. Wash the tissue 3 times with DPBS (add 2% P/S).
3. Use forceps and sterilized scalpels to cut the primary tissue sample on ice into small fragments (approximately < 10cm in size). **!CAUTION** Avoid excessive cutting during the cutting process.
4. Open the grinding chamber, and add the tissue fragments with forceps
5. Run the grinding device, which includes 6 automated steps (a-f): **!CAUTION** Close the cover during operation and be cautious of mechanical hazards.

a. Lower the upper grinding plate until the grinding surfaces fuse together.
b. Add the grinding buffer, set the lower grinding plate rotating at 60 r/min, and start the motor to run for 1 min.
c. Lift the upper grinding plate for 10 mm to separate the grinding plates.
d. Add 10× tissue digestion reagent, and rotate the lower grinding plate for 5min at 60 r/min and 37 °C.
e. Pass the cell suspension through a 100 μm cell filter.
f. Add an equal volume of digestion termination reagent, which contains 10% serum, to the cell suspension.
6. Centrifuge the cell suspension and remove the supernatant. **!CAUTION** When aspirating the supernatant using a pipette, avoid disturbing the cell pellet.
7. Add 2 mL of red blood cell lysis solution to the cell pellet, pipette the mixture, and incubate it at room temperature for 2 min.
8. Re-centrifuge the cell suspension, remove the supernatant, add 2 mL culture medium to the cell pellet, and pipette the mixture to obtain primary cell suspension.
9. Thaw the AO/PI reagent and prepare a premix solution containing 50 µg/mL PI and 10 µg/mL AO in DPBS. **!CAUTION** Avoid contact of AO/PI dye with skin.
10. Resuspend the cells in 1 mL of basal culture medium and mix thoroughly by pipetting up and down.
11. Aspirate 10 µL of the cell suspension and 10 µL of the AO/PI working solution, mix them together, and then add the mixture to a hemocytometer for cell counting. Calculate the cell number.

#### Box 2 | Formulate and print organoid precursor, using the OrgFab printer Timing 1 h Procedure

**▲CRITICAL** The entire process should be placed and operated within a biosafety cabinet to prevent bacterial contamination during use.

1. Centrifuge the cell suspension and remove the supernatant to obtain a cell pellet.
2. Place the cell pellet on ice to pre-chill for 1 min.
3. Use a 100μl pre-cooled pipette tip to aspirate Matrigel on ice, pipette it to the EP tube, and mix with cells by gently pipetting. **▲CRITICAL STEP** The uniformity of the mixture of cells with Matrigel determines the consistency of the organoids. **!CAUTION** During the mixing process, maintain a low temperature at all times and avoid touching the EP tube with hands. **!CAUTION** The Matrigel should not be subjected to repeated freeze-thaw cycles. **!CAUTION** If the matrix gel that has been mixed with cells cannot be used immediately, it should be stored in a refrigerator at 4°C and used within 2-4 hours.
4. Turn on the sterilization switch of the equipment and activate the UV sterilization for 20 min.
5. Install the microfluidic chip to the semiconductor cooling platform, and connect the inlet/outlet ports to corresponding fluids, including two oil inlet ports, one bioink inlet port, and one droplet outlet port. **!CAUTION** Ensure that the chip alignment platform is properly aligned to avoid gaps.
6. Install the collection tubing in the heating chamber, connect the collection port to the droplet formulation outlet port, and connect the other end to the port on the print head. **▲CRITICAL STEP** When connecting the PTFE tubing to the chip component, after securing the tube, confirm whether the tube is in contact with the bottom of the chip. **!CAUTION** During the connection process, do not bend the PTFE tubing.
7. Install the print head, and connect one end of the print head to the collection/incubation tubing and the other end to the high-pressure gas. The third port is placed facing straight to the recipient substrate, e.g. 96-well plate. **!CAUTION** Before installation, confirm the orientation of the print head to ensure proper alignment and maintain airtight connections.
8. Install the cell culture plate and the waste liquid collection trough on the printing platform on the right side of the device. **!CAUTION** Place the culture plate on the plate tray and secure the plate to the tray by aligning and locking it into position.
9. Place the EP tube with bioink in the sample tank and insert the bioink tubing. **!CAUTION** During the installation process, handle the cap of the EP tube and avoid contact with the part of the tube containing the bioink.
10. Initiating OrgFab printer, which includes 8 automated steps (a-h): **!CAUTION** Before starting, ensure that the temperature displayed on the software is set to 4°C. **!CAUTION** Close the chamber lid when starting the printing process and pay attention to mechanical safety. **!CAUTION** During the operation of the equipment, ensure that the ventilation function of the biosafety cabinet is activated. The temperature should be maintained between 20-25°C.

a. Move down the mechanical turntable to fix the chip on the semiconductor cooling platform,
b. Insert the chip bioink inlet tubing into the EP tube for waste collection, and pump oil into the chip channels with flow rates of 312.5uL/min via the two oil inlets to repel air in the channel. Afterward, the chip channels are filled with oil. All fluids are conducted via PTFE tubing. **!CAUTION** Nove7000 is volatile. After adding it, the cap should be closed promptly.
c. Insert bioink inlet tubing into the bioink EP tube, aspirate oil out of the channel from the front right (Oil inlet 1) to port aspirate bioink with flow rates of 250uL/min from the tube, and infuse the channel.
d. Rotate the mechanical turntable (knob) to change the chip channel tandem, which connects the bioink channel to the front left port for the Bioink outlet.
e. Pump oil via the two oil inlets, with flow rates of 312.5uL/min in Oil inlet 2 and 25uL/min in Oil inlet 1.
f. Shear bioink flow into uniform droplets by oil co-flow, forming structural templates of organoid precursors (or cellular spheroids). **!CAUTION** During the initial microsphere preparation, observe the morphology and uniformity of the microspheres. If any abnormalities are detected, click the interrupt button to stop the process and adjust the ratio of the oil-pushing speed.
g. Stop pumping oil toward the two inlet channels, store the cell-laden droplets in the collection tubing, and solidify them at 37°C for 30 min.
h. Pump oil from Oil inlet 1 at 12.5uL/min and Oil inlet 2 at 25uL/min, use a camera and visual detection algorithm to control the instantaneous opening of the high-pressure gas toward the print head. Each air beam firing enables one shot of gelled droplet, i.e. organoid precursor/cellular spheroid, into the 96-well plate, with each well containing one or a certain number of spheroid.
11. Culture organoid precursors/cellular spheroids in a 96-well plate.
12. Disconnect the tubing connected to the chip, remove the chip and tubing, and discard them. **!CAUTION** The chip is a disposable consumable and is classified as biohazardous waste. **!CAUTION** The equipment requires regular maintenance, including monthly disinfection and cleaning of the tubing. **!CAUTION** After wiping the equipment with a lint-free cloth soaked in alcohol, it should be allowed to dry completely before use. Operating the equipment while it is still wet poses a risk of electric shock. **!CAUTION** When cleaning the equipment, appropriate personal protective equipment (PPE) should be worn to prevent adverse effects on the environment and personnel health. **!CAUTION** Do not wipe the internal electrical connections, various ports of the equipment, and the AC power sockets. **!CAUTION** Do not sterilize the equipment using high-temperature, high-pressure, or gaseous disinfection methods.

#### Box 3 | Organoid culture Timing 7-14 d Procedure

1. Open the organoid printer cover and take out the organoid printed on the 96-well plate.
2. Place the plate under a microscope and observe the morphology of the organoid to check the morphology of the organoid and make sure each well of the plate has 1 organoid.
3. Place the culture plate in a 37°C CO_2_ incubator for culture. **!CAUTION** Pay attention to the humidity and CO_2_ concentration of the incubator, and avoid vibrating the plate.
4. Change the medium every other day, until organoids fully grow for 7-14 d.
5. Remove 50 ml of culture medium each time, and then add new culture medium to 100 ml. **!CAUTION** Be careful not to suck up the organoids when changing the culture medium. Tilting the 96-well plate will make it easier to observe.

#### Box 4 | Drug screening Timing 6 d

Procedure

▲**CRITICAL** Choose a light-protected plate when culturing the organoids.

1. For the drug to be tested, prepare a 50 mM drug stock solution and filter it through a membrane. Aliquot 5 μL per tube.
2. After culturing for 7-14 days (depending on the organoid type), place the culture plate under the microscope to observe the organoid structure.
3. Dilute the drug to the highest concentration and then perform a 5-fold gradient dilution. Calculate the volume of drug and culture medium required for each gradient.
4. Use a 1000 μL pipette to add the culture medium in a 15 ml centrifuge tube, and then add the drug stock solution.
5. Use a 100 μL pipette to aspirate and remove the culture plate medium completely.
6. Add the drug-containing culture medium to the plate wells in ascending order of drug concentration. **!CAUTION** For each concentration, set up three replicate wells, and also reserve one additional well for live/dead staining at each concentration.
7. Replace the culture medium with a drug-containing medium to prevent drug degradation every other day.
8. After culture 5 days, thaw the CTG reagent at room temperature and equilibrate the CTG reagent. **!CAUTION** Do not vortex the reagents.
9. After culture 6 days, Thaw the Calcein-AM/PI reagent. In a light-protected EP tube, sequentially add 900 μL ddH₂O, 100 μL 10× Assay Buffer, 1 μL Calcein-AM Solution (2 mM), and 3 μL PI Solution (1.5 mM). Mix them and set aside for use.
10. Using a 100 μL pipette, aspirate the culture medium from the live/dead staining wells. Wash with DPBS, then add the Calcein-AM/PI working solution.
11. Stain at room temperature in the dark for 5 minutes. Then, using a 100 μL pipette, aspirate the staining solution, wash with DPBS, and observe under a fluorescence microscope for imaging.
12. Add 100 μL of CTG reagent to each well (maintaining consistency with the culture medium volume). Place the culture plate on a shaker and shake for 2 minutes. Use a microscope to observe whether the organoid spheroids have dispersed. If not dispersed, use a multichannel pipette to pipette up and down 10-15 times, and then recheck under the microscope to ensure the spheroids are dispersed. **!CAUTION** When using a multichannel pipette to blow and mix, be careful to avoid creating bubbles.
13. Incubate at room temperature for 10 minutes. Record the luminescence (LUM) signal intensity using a microplate reader and construct an IC50 curve.

#### Box 5 | Laminate organoids on the chip Timing 30 min

Procedure

1. Place the cell culture plate under the microscope.
2. Select and pick out organoids using a Pasteur pipette under the guidance of a microscope.
3. Aspirate the culture medium from the wells using a pipette and wash with 1× SSC buffer. **!CAUTION** When using the pipette, take care not to touch the organoids.
4. Add cell recovery solution for the organoid and incubate it at 4°C for 15min
5. Wash the cell recovery solution for organoid with 1× SSC buffer, add cell dissociation reagent, and incubate at room temperature for 2 min.
6. Place a drop of ddH₂O on the microscope slide and adhere the chip to the slide. **!CAUTION** If lamination is performed on the microscope slide, this step is not required.
7. Use a Pasteur pipette to load the organoid on the chip/microscope slide. **!CAUTION** When using a Pasteur pipette, take care not to touch the surface of the chip.
8. Use a gas cylinder to blow and dry the liquid around the organoid. **▲CRITICAL STEP** Remove the liquid to ensure that there is no slipping during the lamination process. **!CAUTION** When using a gas cylinder, pay attention to the gas flow rate to avoid blowing away the organoids. **!CAUTION** The surface of the organoids needs to be dried as much as possible, but complete desiccation should be avoided. A thin layer of water film should be retained on the surface of the organoids.
9. Use scissors to cut the PTFE film into a size of 3 cm × 3 cm. **!CAUTION** During the cutting, avoid the creased sections of the PTFE film.
10. Use tweezers to cover the chip with a piece of PTFE film. **!CAUTION** Avoid the formation of wrinkles during the covering process. If the cover is misaligned, do not lift it to reposition. **!CAUTION** This step should be completed as quickly as possible to avoid the disappearance of the water film on the organoids.
11. Install the microscope slide on the lamination device.
12. Place another glass slide on the lamination device and place a weight on top of the glass slide **!CAUTION** When installing another microscope slide, align the edge of the slide with the raised platform on the device. **!CAUTION** Before installing the glass slide, lower the manual height adjustment stage to its lowest position to avoid contact between the glass slide and the PTFE film.
13. Switch the knob to lift the chip until it touches the glass slide installed on the top. **!CAUTION** The upper microscope slide should be detached from the boss and maintained in this position for 10s.
14. Remove the weight and the glass slide gently and vertically in sequence. **▲CRITICAL STEP** When removing the weights and the microscope slide, avoid any movement or sliding of the PTFE film.
15. Uninstall the chip from the lamination device, heat the chip at 37°C for 5min, and use tweezers to remove the PTFE film. **▲CRITICAL STEP** Use tweezers to lift the PTFE film from one side, ensuring it does not slip.
16. Immunofluorescence staining of a laminated organoid, showing DAPI staining of the laminated cellular spheroid.
17. Laminated organoids on a spatial transcriptomic chip, enabling sequencing and resolution of cell types and cellular interactions.

## Materials

### Biological Materials

#### Ethics statement

Human samples were collected after informed written consent was obtained from all donors in accordance with study protocols conforming to the provisions of the Declaration of Helsinki. For the pediatric participants, written informed consent was obtained from adult legal guardians of the children before enrolment. The study was approved by the Ethics Committees of Tsinghua Shenzhen International Graduate School (Refs. 202171 and 202295).

#### Mouse

Male BALB/c mice (4-6 weeks old) were purchased from the Medical Laboratory Animal Center of Guangdong, China, and housed in the animal facility of Tsinghua University Shenzhen International Graduate School (SIGS). Environmental conditions were maintained at a temperature of 23±2°C, with a relative humidity of 50-65%, in a pathogen-free environment, under a 12-hour light/12-hour dark cycle, with *ad libitum* access to food and water.

#### Human

Patient-derived colorectal cancer and liver cancer samples were obtained from Shenzhen People’s Hospital, Shenzhen, China (Ethical Development No. 2024-08401).

**!CAUTION** The use of human tissues and human stem cells must adhere to institutional and funding body regulations, as well as relevant ethical guidelines.

### Reagents

#### Tissue digestion solution

- Collagenase Ⅰ (**Y**uanye, cat. no.S10053-1g)
- Collagenase Ⅳ (**Y**uanye, cat. no. S10056-1g)
- Dispase Ⅱ (**Y**uanye, S25046-1g)
- DNase (Solarbio, D8071)
- DPBS (Biosharp, cat. no.BL310A)
- FBS (QmSuero, mu001SR)
- P/S (Biosharp, BL505A)
- AO/PI (Countstar, cat. no. RE010212)

#### Organoid formulation and culture

- Engineered Fluid (3M Novec, cat. no.25556)
- MasterAim® organoid complete medium for colorectal cancer (MasterAim, cat. no.10-100-018)
- MasterAim® organoid complete medium for liver cancer (MasterAim, cat. no.10-100-296)

#### Organoid lamination and characterization

- Cell Recovery Solution (Corning, cat. no.354253)
- Methanol (Rhawn, cat. no.R007536)
- Trypsin (Gibco, cat. no.15050057)
- PTFE (Millipore, cat. no.IPVH00010)
- BSA (for non-sterile use) (Biosharp, cat. no. BS114-100g)
- DAPI (Biosharp, cat. no.BL739A) **!CAUTION** Due to DAPI’s photosensitivity, avoid exposing it to light.
- AntiFade Mounting Medium (Beyotime, cat. no.P0131)
- 4% Paraformaldehyde (PFA) (Biosharp, cat. no. BL539A) **!CAUTION** Paraformaldehyde (PFA) is toxic and irritating. Handle with care.
- PBS (nonsterile) (Biosharp, cat. no. BL601A)
- Sucrose (Solarbio, cat. no. S8271)
- Tissue-Tek O.C.T. compound (Biosharp, cat. no. BL557A)
- TritonX-100 (Solarbio, cat. no. T8200)
- Tween20 (Leagene, cat. no. PW0026)
- CEA (Huabio, cat. no.ER1906-09)
- Lysozyme (Huabio, cat. no.ET1609-35)
- MUC1 (Huabio, cat. no. ET1611-14)
- FAPB1 (Huabio, cat. no.ET1704-23)
- RDX (Huabio, cat. no. ET1610-41)
- AFP (Huabio, cat. no. EM1701-31)
- OPN (Huabio, cat. no. EM1701-31)

#### Drug screening

- CellTiter-Glo®Luminescent Cell Viability Assay (Promega, cat. no.G7570)
- Calcein-AM/PI (Solarbio, cat. no.CA1630)
- CTG reagent (Promega, cat. no. G7572)
- 5-FU (Aladdin, cat. no. F100149-1g)
- Epirubicin (Abmole, cat. no. M5617)
- Oxaliplatin (MCE, cat. no. HY-17371)
- Gefitinib (Aladdin, cat. no. G125799)
- Gemcitabine (Aladdin, cat. no. G127944-5g)
- Cisplatin (Aladdin, cat. no. HY-17394)
- Paclitaxel (Aladdin, cat. no. P106869)
- Anti-PD-1 (Aladdin, cat. no. C412007)
- Lenvatinib (Aladdin, cat. no. L125518)

### Reagent Setup

#### 10× Tissue Digestion Reagent

Take 200 mg Collagenase IV, 200 mg Collagenase III, 200 mg Collagenase I, 200 mg Dispase, and 13.33 mg DNase, dissolve them in 10 ml DPBS, vortex to dissolve, and after ensuring complete dissolution, filter through 0.22 μm filter to sterilize.

**!CAUTION** It is necessary to pass the cell filter through the filter slowly and keep the reagent drops being deposited as droplets when executing, to avoid damage to the filter.

#### Digestion Termination Reagent

Take 9 ml of DMEM/F12, 1 ml of FBS, and 100 μl of P/S, and mix by pipetting with a 1000-μl pipette.

#### CTG reagent

Thaw the CTG buffer at room temperature. Equilibrate the CTG substrate dry powder to room temperature. Mix the CTG buffer and CTG substrate in a light-protected centrifuge tube, and gently vortex to mix thoroughly, forming the CTG reagent. Aliquot 1000 µL per light-protected EP tube and store at −20°C.

### Equipment

- P10 micropipette (Eppendorf, cat. no.3124000040)
- P100 micropipette (Eppendorf, cat. no.3124000075)
- P1000 micropipette (Eppendorf, cat. no.3124000121)
- P10 pipette tips (Cellprobio, cat. no.800301)
- P100 pipette tips (Cellprobio, cat. no.801101)
- P1000 pipette tips (Cellprobio, cat. no.FT-1250)
- Biosafety cabinet (ESCO, model. no.AB2-S-CN)

#### Tissue digestion solution

- Disposable scalpel (Kyuan, cat. no.0408)
- Petri dishes, 60 mm (Thermo, cat. no.)
- Tweezers, stainless steel (Thermo, cat. no.DS0399-0001)
- Cell strainers, 100 μm (Biologix, cat. no.15-1100)
- Countstar Mira FL (Countstar, model. no.P020300022)
- Orgfab® cell extractor (Synorg, model. no.)

#### Organoid formulation and culture

- rgfab**^®^** Organoid 3D printer (<u>Synorg, model. no.P01</u>)^54,55^ For manual microfluidic-based formulation of droplet organoid precursors, see Supplementary Figure 1
- tubing (Woer, cat. no.24T)
- incubator (ESCO, model no.CLM-170B-8-TC)

#### Organoid lamination and characterization

Organoid lamination device

Freezing microtome (Leica, model. no.CM1860)

Fluorescence microscope (Nikon, model. no. ECLIPSE Ts2-FL)

### Software

- NIS-Elements (Nikon microsystems, provided with the microscopes)
- ImageJ (ImageJ-win64)
- GraphPad (GraphPad.Prism.9.0.0.121)

## Troubleshooting

Troubleshooting advice can be found in Table 1.

**Table 1.** Troubleshooting table.

| Table 1 \| Troubleshooting table |  |  |  |
| --- | --- | --- | --- |
| Step | Problem | Possible reason | Solution |
| Stage 1.5a | More tissue remains in the filter | The grinding surface does not fit in the grinding mode | Check the grinding surface fit height |
| Stage 1.5a | Low viability of the extracted cells | The sample tissue is more fragile | Check grinding pressure and reduce grinding time |
| Stage 1.5d | Fewer cells extracted | Tissue digestion enzymes are not at working temperature | Check the equipment temperature record and heating module, and extend the digestion time |
| Stage 1.7 | The extracted cells contain more red blood cells | Insufficient time for treatment with red blood cell lysis buffer | Resuspend the cell pellet with red blood cell lysis buffer and let it stand for 2 minutes. |
| Stage 2.3 | Bubbles in bioink | Bubbles generated when preparing bioink | Use a centrifuge to remove air bubbles and remix |
| Stage 2.3 | Bioink is not sheared into droplets | Hands touched during the preparation of bioink | Re-prepare the bioink and handle it with caution. |
| Stage 2.5 | Bubbles in the fluorine oil of the microfluidic chip | The pipe is not connected properly | Check pipe connections |
| Stage 2.8 | Print head cannot align with the orifice plate | The culture plate is not aligned with the slot | Realign the slots to install the culture plate |
| Stage 2.10.f | Bioink is not sheared into droplets | High ambient temperature, high freezing temperature | Check process temperature control and ambient temperature |
| Stage 2.10.f | Organoid precursor printing is unstable | Gas in the tubing | Check tubing connections |
| Stage 2.10.h | Printing nozzle identifies organoids incorrectly | Insufficient light source in the nozzle | Check the nozzle light source |
| Stage 2.10.h | No organoid precursors in printing | The tubing is squeezed or folded during installation | Check the installed tubing |
| Stage 3.2 | Organoids adhere to the culture | Not using low attachment culture plates | Use low-adhesion culture plates and increase shaker speed |
| Stage 3.4 | Medium yellow/acidic before the next feeding | The culture medium evaporates too quickly | Check the incubator tank and add extra culture medium to the cell culture plates |
| Stage 3.4 | Organoids disintegrate in culture | Too many cells in organoids | Check the cell count in Stage 1 and modify the cell and Matrigel mix ratio |
| Stage 4.13 | A large difference in IC50 data among the test replicates. | Air bubbles in the culture plate | Check under a microscope for bubbles |
|  |  | The original medium was not drained when adding the medium containing the drug | Pay attention to the pipetting operation, change the small-range pipette tip for pipetting |
| Stage 5.5 | The organoids are broken after dissociation | Different types of organoids have different densities and require specific processing times | Reduce incubation time depending on the organoid type |
| Stage 5.7 | Losing organoids when transferring | Repeated pipetting causes organoids to break apart | Avoid repeated blowing |
|  |  | Organoids adhere to the inside of the Pasteur pipette. | Re-aspirate and slowly push the organoid out |
| Stage 5.12 | Uneven lamination between the left and right parts of the organoid | The weight is not placed in the center of the chip | Align the weights with the concentric circle markings on the laminating machine |
|  | Organoids still have cell stacks after lamination | Different organoids have different density | Increase weight pressure |
| Stage 5.14 | Laminated organoids show tailing | No vertical removal of weights and slides | Be careful not to damage the PTFE film when removing the weights and glass slide. |
| Stage 5.15 | Organoids adhered to the PTFE film | More liquid is left in Stage 5.8 | Use a gas cylinder to remove as much excess liquid as possible |
|  | Laminated organoids show tailing | Slippage occurs when removing the PTFE film | Use pointed tweezers to peel off the PTFE film from one corner |

## Timing

Step 1, cell extraction, 30 min

Step 2, organoid formulation and printing, 1 h Step 3, organoid culture, 7-14 days

Step 4, drug screening, 6 days

Step 5, organoids lamination and characterization, 20 min (lamination) + 2 days (Immunostaining)

## Anticipated Results

This protocol establishes an automated workflow for the construction and characterization of patient-derived primary tissue ex vivo models, aiding in therapeutic and mechanistic studies. The method was primarily applied to tissues obtained from patient surgeries and biopsy sampling. We applied the protocol to a surgically resected specimen of colorectal cancer (Figure 5A) and liver cancer (Figure 6A).

**Figure 5.**
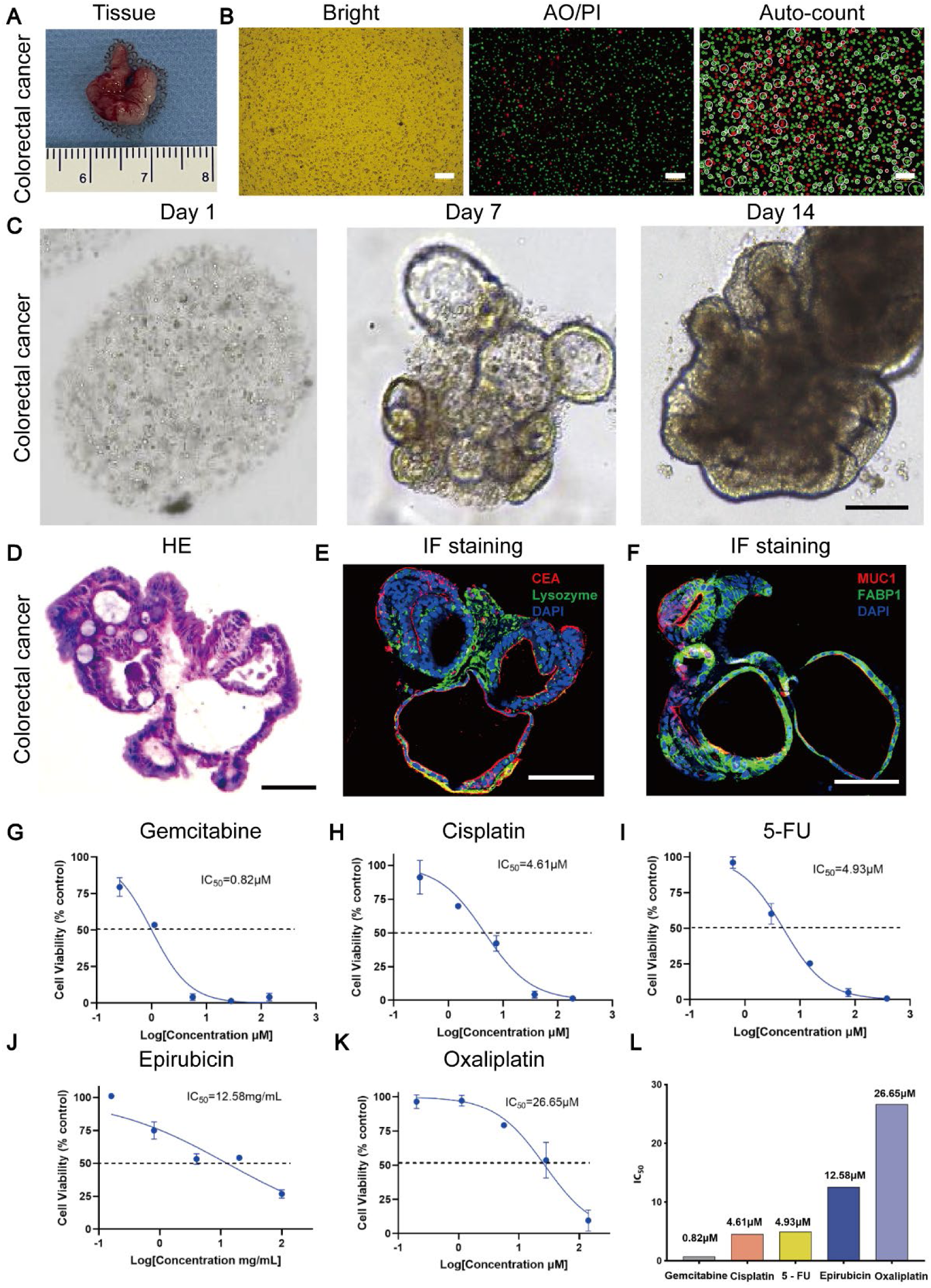
Results of device-extracted colorectal cancer primary cells and formulated DEOs. A) Colorectal cancer tissue B) Bright-field, live/dead staining, and cell count images of primary cells extracted after Step 1, Scale bar, 100μm C) Bright-field image of cellular spheroids of Step 2, Scale bar, 10μm D) HE staining of colorectal cancer organoid slices, Scale bar, 10μm E-F) Immunofluorescence staining of colorectal cancer organoid slices. Figure E, red: CEA, green: Lysozyme, blue: DAPI. Figure F, red: MUC1, green: FABP1, blue: DAPI. Scale bar, 100μm G-K) IC50 test curves and statistical graphs of colorectal cancer organoids in Gemcitabine, Cisplatin, 5-FU, Epirubicin, and Oxaliplatin drugs. The results show that Gemcitabine has the best inhibitory effect on the patient’s colorectal cancer organoids

**Figure 6.**
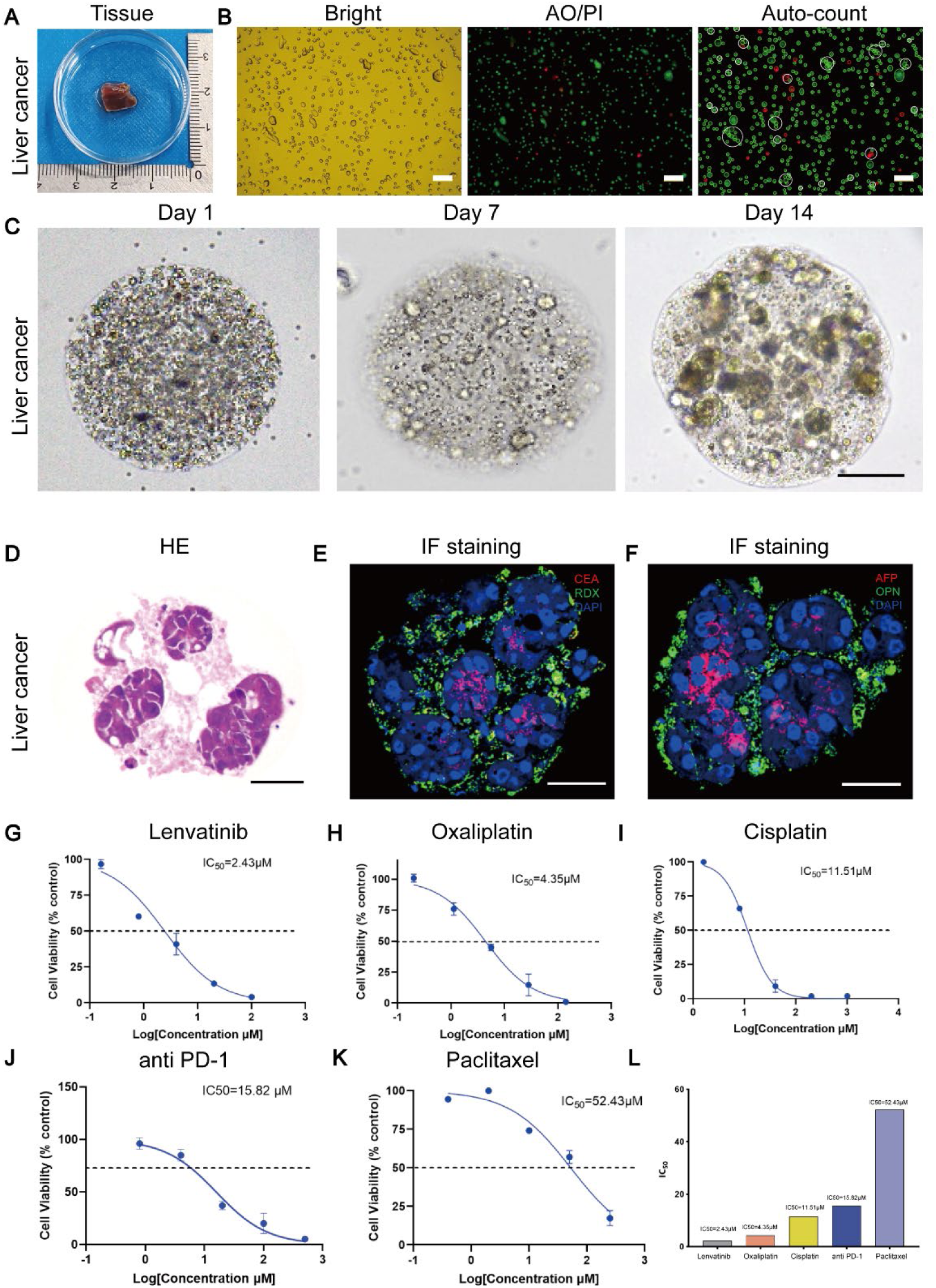
Results of device-extracted liver cancer primary cells and formulated DEOs. A) Liver cancer tissue B) Bright-field, live/dead staining, and cell count images of primary cells extracted after Step 1, Scale bar, 100μm C) Bright-field image of cellular spheroids of Step 2, Scale bar, 10μm D) HE staining of liver cancer organoid slices, Scale bar, 10μm E-F) Immunofluorescence staining of liver cancer organoid slices. Figure E, red: CEA, green: RDX, blue: DAPI. Figure F, red: AFP, green: OPN, blue: DAPI. Scale bar, 100μm G-K) IC50 test curves and statistical graphs of liver cancer organoids in Lenvatinib, Oxaliplatin, Cisplatin, Anti-PD-1, and Paclitaxel drugs. The results show that Lenvatinib has the best inhibitory effect on the patient’s colorectal cancer organoids

Stage 1: Extraction and Purification of Primary Cells from Tissue Samples

In Stage 1, the automated cell extraction device, sTED, processed ultra-small tissue samples, such as the surgically resected colorectal cancer tissue (Figure 5A) and liver cancer tissue (Figure 6A). The device employed a tissue grinding strategy with custom-designed, disposable grinding plates, enabling cell extraction from tissue fragments less than 1 mm in size. After using the cell extraction device, we observed a higher number of cells with greater viability (Figure 5B, Figure 6B). Viability assays using AO/PI staining and automated cell counters facilitated accurate assessment of cell viability and yield. Bright-field microscopy provided visual confirmation of the extracted cells, showing a high number of viable cells with minimal debris. Cell viability rates exceeded 60-90% for the patient tissues, depending on the inherent heterogeneity and necrotic regions commonly attributed to malignant tissues. The entire cell extraction process was completed within approximately 30 minutes, an important factor when working with ultra-small samples.

Stage 2: Formulation and Bioprinting of Cell-Laden Microspheres

In Stage 2, we utilized the OrgFab integrated bioprinter to rapidly produce uniform cell-laden microspheres (organoid precursors). After mixing the extracted primary cells with Matrigel to create the bioink, the mixture was processed through microfluidics to generate uniform droplets via oil-phase shearing. The printed organoid precursors exhibited high uniformity in size and morphology, as demonstrated by homogenization violin plots of organoid size, shape, and cell density (Figure 3H, Supplementary Figure 2A). The tight distributions indicated consistent fabrication across wells, and images of the organoid precursors in a 96-well plate (Figure 5C, Figure 6C) showed uniform spheroids suitable for downstream applications. A bioink volume of 10 μL produced approximately 100 organoid precursors (spheres), sufficient for a batch of drug testing. The OrgFab printer operated in a fully automated manner, handling ultra-small samples down to 10 μL, from sample injection to organoid patterning. The formulation and printing process was completed within 1 hour, enabling efficient preparation of organoids for culture and experimentation. Post-printing, bright-field microscopy confirmed the morphology and uniformity of organoid precursors (Supplementary Figure 2). The high-density microfluidic droplet encapsulation promoted consistent microenvironment recovery, facilitating organoid maturation during the subsequent culture period (Figure 5C, Figure 6C). Over a 14-day culture period, we observed that as the organoids grew, hollow structures gradually formed within them, indicative of lumen formation and the development of tissue-like architecture.

Stage 3: Lamination of Organoids for Rapid Characterization

In Stage 3, the cultured organoids were ready to be laminated into a monolayer using the lamination device. Based on the dye diffusion rate characteristics, we used DiI dye to rapidly mark the organoids before lamination, selectively staining only the outer cells of the organoids’ internal structures^56^. After lamination, we utilized ss-DNA to label the cell nuclei. By comparing the differential expression data, we identified corresponding cell clusters in both 2D and 3D configurations (Figures 7D and 7E), and we observed clear cell positions, with each cell distinctly separated (Figure 7).

**Figure 7.**
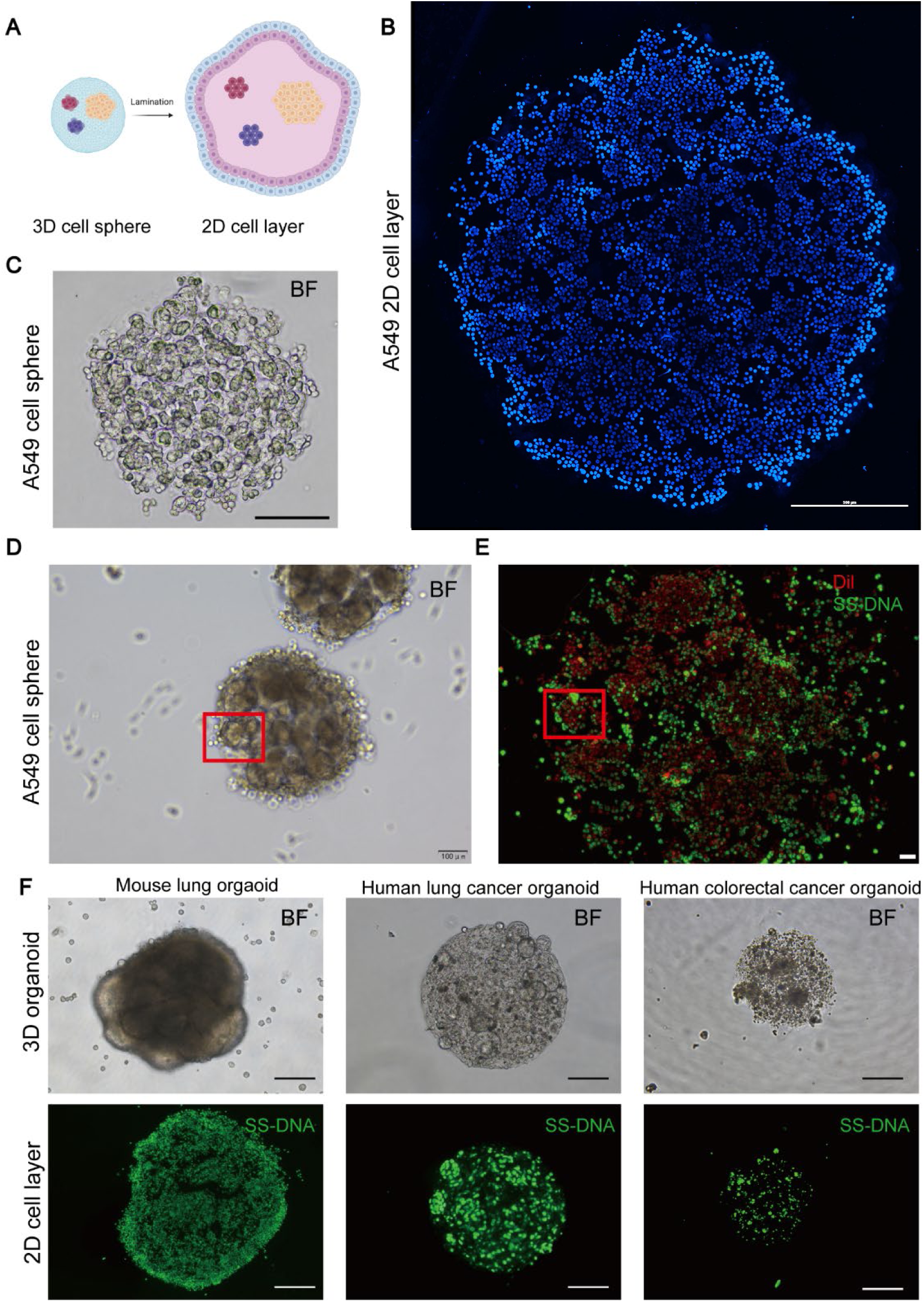
Results of organoid lamination. A) Schematic image of lamination B) Bright-field (10x) of 3D A549 cell sphere, Scale bar, 100μm C) DAPI immunofluorescence staining images of cell spheres of B, Scale bar, 500μm D) Bright-field (4x) of 3D A549 cell sphere, Scale bar, 100μm E) Immunofluorescence staining of liver cancer organoid slices. red: Dil dye, green: ss-DNA F) Lamination of different organoid

It is possible to characterize laminated samples through H&E, immunofluorescence staining, and spatial transcriptomic profiling. We performed H&E staining on the colorectal and liver cancer organoids to verify their growth and the retention of primary cell characteristics (Figure 5D, Figure 6D, Supplementary Figure 3).

Immunofluorescence staining of the laminated organoids revealed well-preserved cellular morphology and distribution (Figure 4E), confirming that the lamination compressed the organoids into a single-cell-layer morphology without disrupting cellular architecture and epithelial structures, allowing for comprehensive analysis. Staining with antibodies against specific markers for colorectal cancer (CEA, Lysozyme, MUC1, and FABP1) and liver cancer (CEA, RDX, AFP, OPN) indicated the presence of diverse cell types within the organoids (Figures 5E, 5F and Figures 6E, 6F). The varying expression levels of these markers validated the differentiation and cellular heterogeneity of the organoids, mirroring the complexity of native tissue.

Laminated organoids placed on a spatial transcriptomic chip (Figure 4F) allowed for high-resolution mapping of gene expression. This technique resolved various cell types within the organoids (Supplementary Figure 3), providing insights into cellular interactions and tissue organization. Furthermore, confocal fluorescence microscopy and spatial transcriptomics offered detailed information on cellular heterogeneity and tumor microenvironment interactions.

Applications: Drug Screening and Personalized Medicine

The protocol’s effectiveness was tested through functional applications, here specifically for drug screening using patient-derived organoids from colorectal and liver cancers. For the colorectal cancer organoids, we evaluated five chemotherapeutic agents: Gemcitabine, Cisplatin, 5-Fluorouracil (5-FU), Epirubicin, and Oxaliplatin, and plotted the IC50 curves (Figures 5G-K). Gemcitabine exhibited the lowest IC50 value, demonstrating the most potent inhibitory effect on the patient’s organoids. Based on these findings, we suggested Gemcitabine as the most effective chemotherapeutic option for the patient (Figure 5L).

Similarly, for the liver cancer organoids, we tested the multikinase inhibitor Lenvatinib; the chemotherapeutic agents Oxaliplatin, Cisplatin, and Paclitaxel; and the Anti-PD-1 antibody, generating IC50 curves for each drug (Figures 6G-K). Lenvatinib showed the lowest IC50 value, indicating the most potent inhibitory effect on the patient’s liver cancer organoids. These results confirm Lenvatinib as a potentially effective treatment option for the patient (Figure 6L). Both cases demonstrate the protocol’s applicability in personalized medicine.

Cell viability assays, such as the CellTiter-Glo luminescent assay, quantified the organoids’ responses to drug treatments. Live/dead staining with Calcein-AM/PI provided visual confirmation of cell viability, enabling the assessment of drug efficacy. This direct application of organoid-based drug screening to a clinical scenario is used to demonstrate the translational potential of the protocol.

## ACKNOWLEDGEMENT

The work was supported by the National Key-Area Research and Development Program of China (2024YFA0919800); National Natural Science Foundation of China (32371470 and 82341019); Merck Research Grant, and the Cross-disciplinary Research and Innovation Fund of Tsinghua SIGS (No. JC2022007); Guangdong Basic and Applied Basic Research Foundation (2023B1515120025); Shenzhen Fundamental Research Program (No. JCYJ20240813112004006); Shenzhen Major Science and Technology Projects (KJZD20230923115400001).

## CONTRIBUTIONS

S.M., Y.C., and X.D. designed the study. W.W., Y.H., L.J. and Y.Z. performed the experiments and analyzed the data. W.W. and D.K. wrote the manuscript. J.W. and X.Y. performed the immunofluorescence staining and animal experiments. S.M. supervised the work and revised the manuscript. All authors read and approved the final manuscript.

## DATA AVAILABILITY

The data presented below the figure panels are available from the original references listed in the figure legends. All other data were generated for this article. Any raw data are available for research purposes from the corresponding author upon request. Devices used in this paper are also available from the corresponding author upon request.

## Supplementary Figures

**Supplementary Figure 1.**
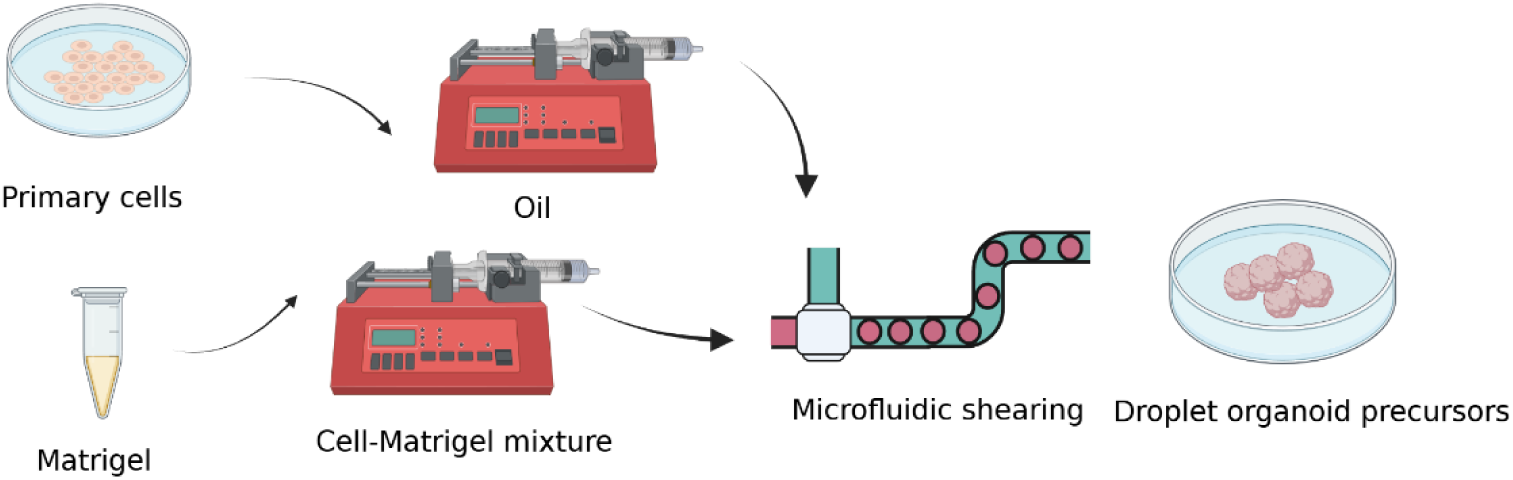
Manual fabrication of microfluidic channels for generating droplet organoid precursors. Timing 2 d Procedure ▲ **CRITICAL** This entire process must be performed in a sterile environment (e.g., biosafety cabinet or cleanroom) to prevent bacterial contamination.

1. Combine polydimethylsiloxane (PDMS) base and curing agent (Corning, cat. no. 04019826) at a 10:1 ratio in a clean container. Stir thoroughly using a glass rod or plate for 15 min to ensure homogenous mixing.
2. Pour the mixture into a pre-cleaned molding mold and place it under a vacuum pump to remove air bubbles. ▲ **CRITICAL STEP** The efficiency of vacuum degassing significantly affects the performance and integrity of the final microfluidic channels. **! CAUTION** Ensure the mold is level and stable to prevent uneven casting.
3. Transfer the mold to a dry oven at 85 °C and cure overnight. Allow the PDMS to cool to room temperature, then cut into ~1 cm³ cubes using a scalpel or blade.
4. Using a 1 mm diameter biopsy punch, make three ports (inlet, outlet, and collection) in each PDMS cube.
5. Connect 24T microfluidic tubing (Woer, cat. no. 24T) to the punched holes in the PDMS block. Attach the inlet to a 1 ml syringe (for cell-Matrigel mixture), the second inlet to a 10 ml syringe (for carrier fluid), and the outlet to a 50 cm segment of 24T tubing to collect the droplet organoid precursors. ▲ **CRITICAL STEP** Confirm all tubing connections are airtight and securely inserted into the PDMS block to maintain flow stability.
6. Load materials:

a. Load the 1 ml syringe with a pre-prepared cell-Matrigel mixture.
b. Load the 10 ml syringe with sterile 7000 Engineered Fluid (3M Novec, cat. no. 25556).
c. Mount both syringes onto a dual-channel syringe pump. ! **CAUTION** Ensure all syringes are free of air bubbles before connection.
7. Configure the syringe pump to the following settings:

a. 1 ml syringe (dispersed phase): 8 µl/min
b. 10 ml syringe (continuous phase): 48 µl/min
8. Start droplet formation:

a. Begin by activating the 10 ml syringe pump to pre-fill the PDMS channels and tubing with carrier fluid.
b. Next, initiate the 1 ml syringe pump to begin generating droplet organoid precursors.
9. After sufficient droplets are collected in the outlet tubing, detach it from the PDMS block and incubate the tubing at 37 °C for 30 min to solidify the Matrigel within the droplets.
10. Connect one end of the collection tubing to a 10 ml syringe filled with PBS. Carefully align the other end with a culture dish containing prewarmed medium. Gently push the PBS through the tubing to dispense the solidified droplet organoid precursors into the dish. **! CAUTION** Avoid high flow rates when expelling droplets, as excessive shear force may deform the droplet structure.

**Supplementary Figure 2.**
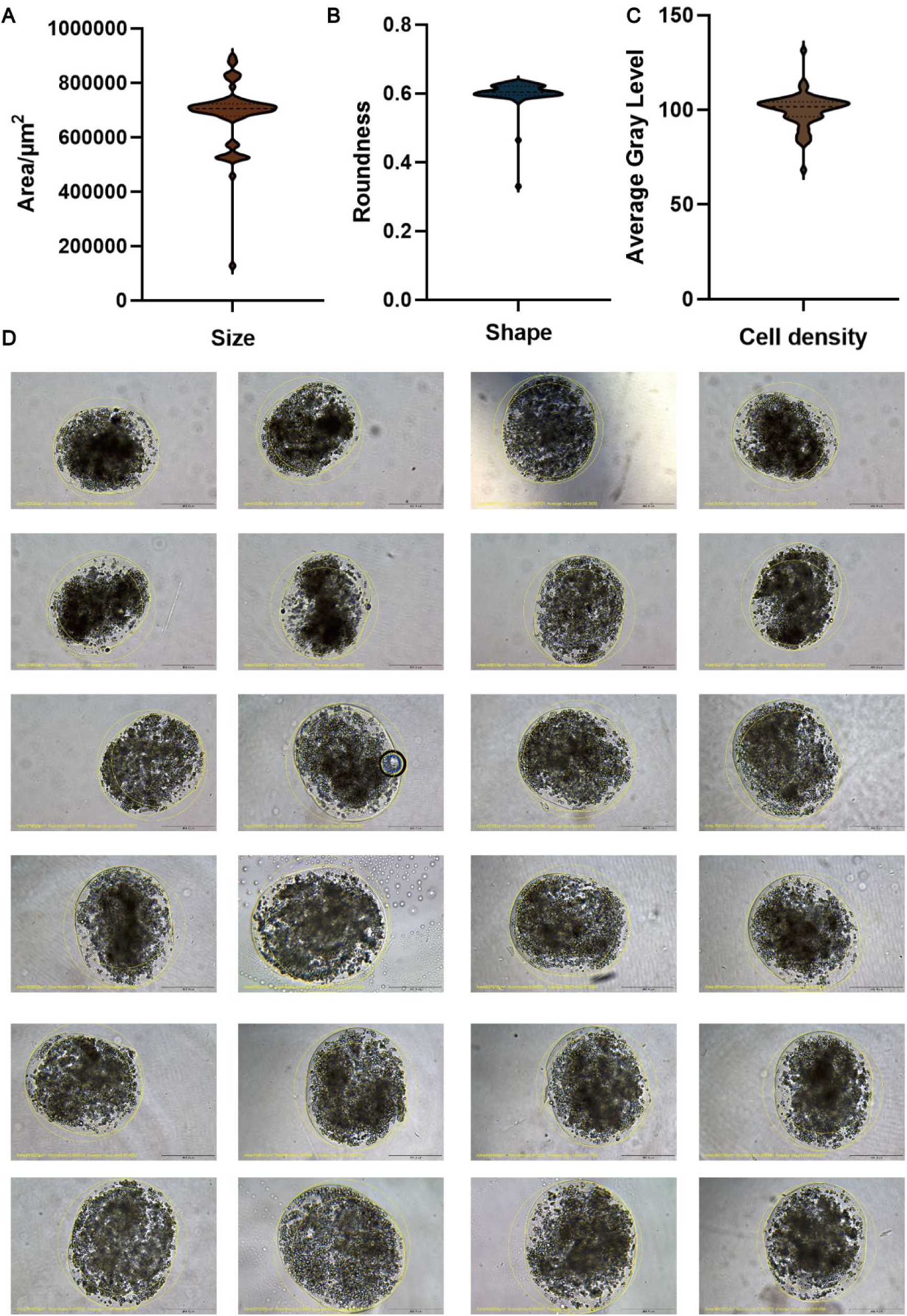
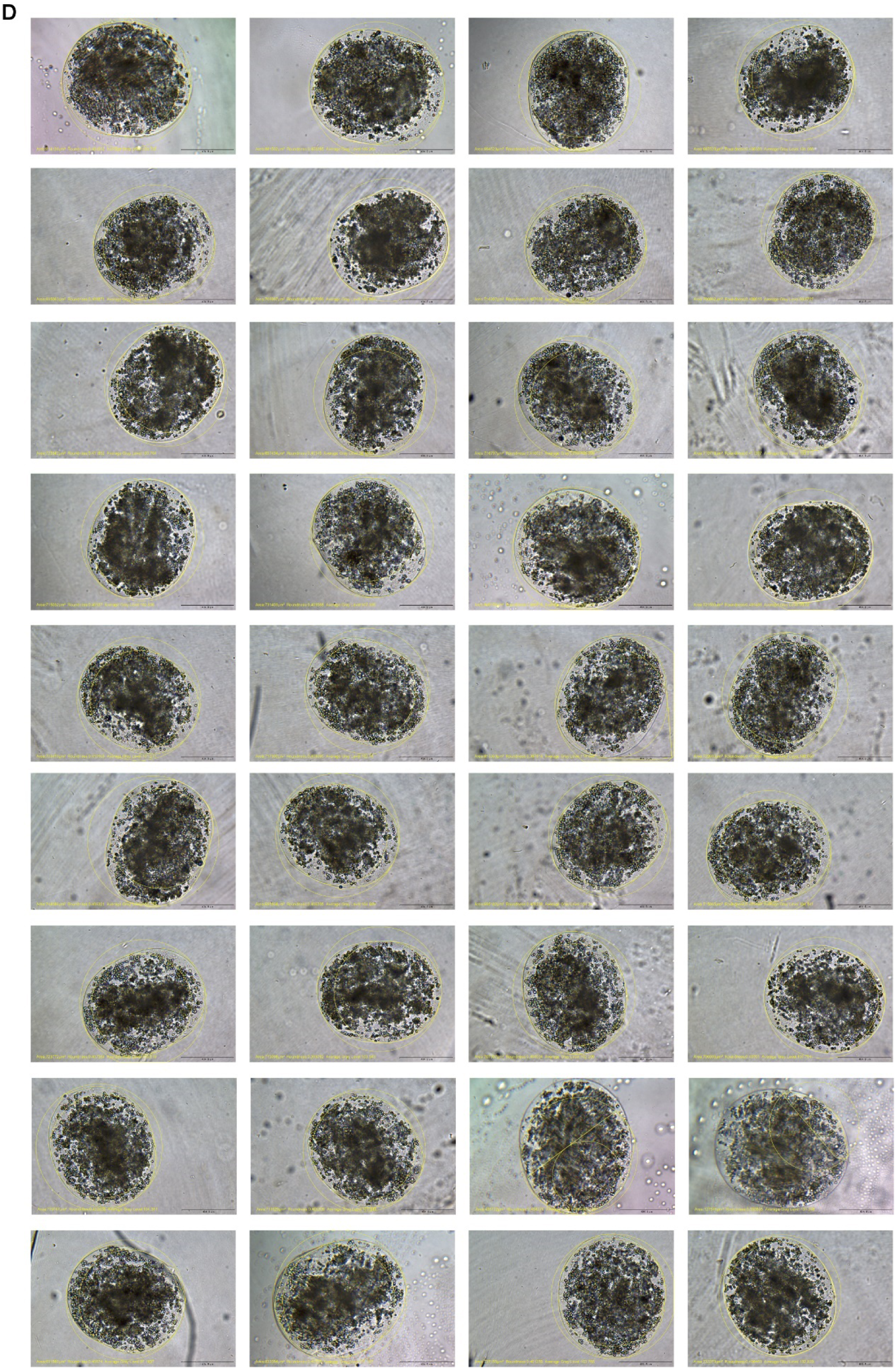
Results of formulating organoid precursor, using the OrgFab printer. A-C) Violin plot of organoid precursor size, shape, and cell density statistics D) Bright-field images of organoid precursors and analysis results

**Supplementary Figure 3.**
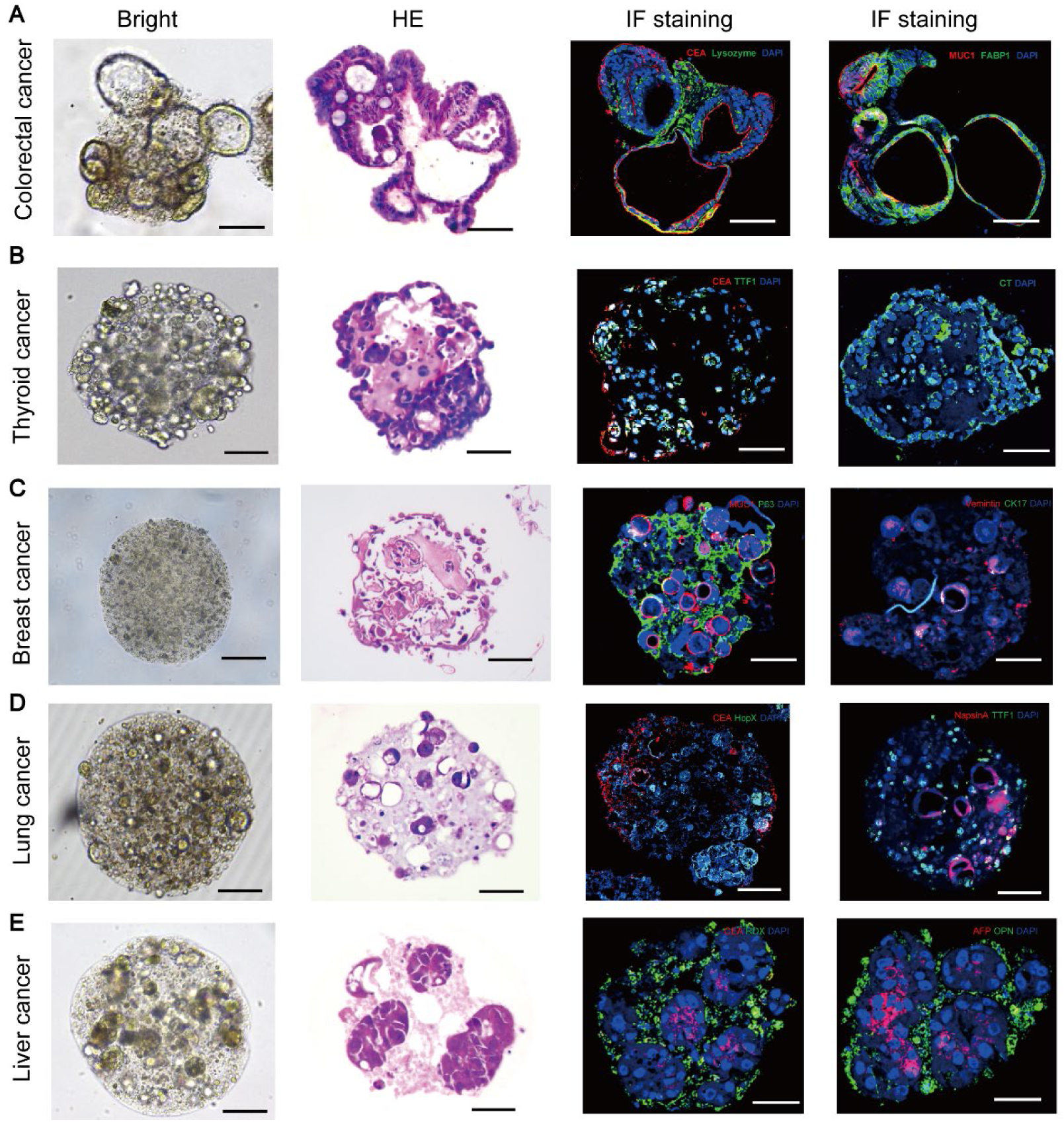
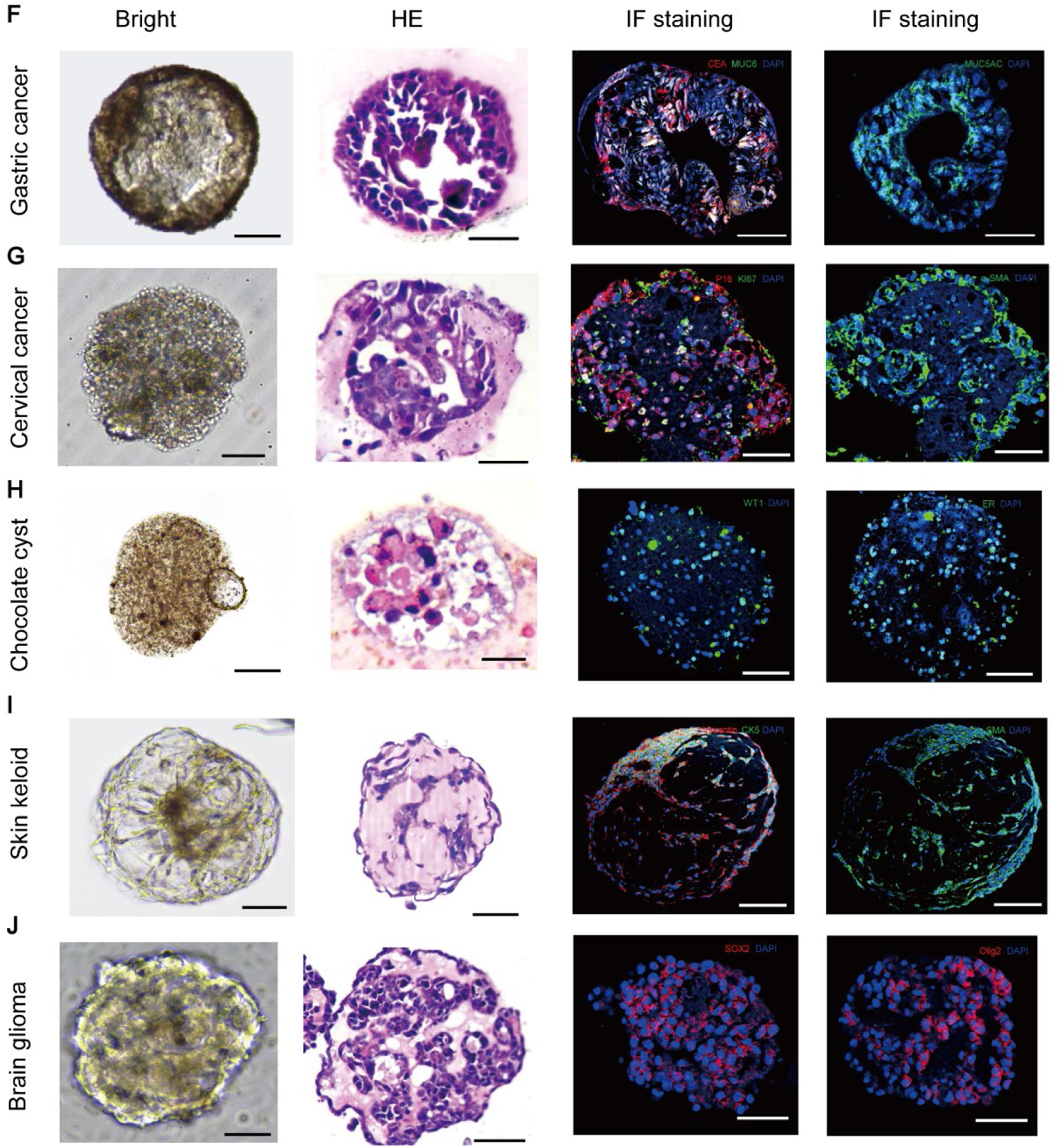
Results of different tissue DEOs. A. Colorectal cancer organoid
B. Thyroid cancer organoid
C. Breast cancer organoid
D. Lung cancer organoid
E. Liver cancer organoid
F. Gastric cancer organoid
G. Cervical cancer organoid
H. Chocolate cyst organoid
I. Skin keloid organoid
J. Mouse brain glioma organoid

